# ER oxidoreductin-1_α_ and unfolded protein response as sex-dependent drivers of cardiorenal dysfunction in experimental autoimmune encephalomyelitis

**DOI:** 10.1101/2025.10.06.680703

**Authors:** Kayla L. Nguyen, Kelly Tammen, Annie McDermott, Sreejita Arnab, Ari Hoffman, Maeva Talla, Nikhil Agarwal, Erin Jones, Aravind Meyyappan, David Mendelowitz, Yair Argon, John R. Bethea

## Abstract

**Background:** Multiple sclerosis (MS) is associated with increased cardiovascular and renal morbidity, but mechanisms linking CNS autoimmunity to peripheral organ injury remain poorly defined. We tested the hypothesis that experimental autoimmune encephalomyelitis (EAE) induces cardiorenal dysfunction via sex-specific dysregulation of endoplasmic reticulum (ER) oxidoreductases and unfolded protein response (UPR) signaling.

**Methods:** Adult female and male C57BL/6J mice [10-12 weeks] and IRE1_α_C148S knock-in mice underwent non-pertussis toxin EAE (nPTX-EAE) induced with 100µg MOG_35-55_ in CFA and 200µg heat-inactivated MTB with a booster at 7 days. Controls received all components except MOG. Motor scoring was done daily and EN460 (ERO1_α_ inhibitor, 10 mg/kg IP) was given twice weekly beginning at 10DPI. Echocardiography and renal Doppler (Vevo 2100) were performed at 36-38DPI with primary outcomes of LV systolic/diastolic function and renal perfusion with tissue collected at 40DPI. LV and kidney were analyzed via western blot for ERO1_α_, PDIA1, Prdx4, 4HNE, and BiP. Data are expressed as mean ± SEM with two-way ANOVA/Tukey’s post hoc or Mann Whitney test performed and outliers identified by ROUT (Q=1%).

**Results:** Sixty-seven percent of immunized mice developed motor deficits. At 36-38DPI, EAE reduced ejection fraction and fractional shortening while increasing IVCT, IVRT, and myocardial performance index without hypertrophy. Renal resistive index increased and end-diastolic velocity decreased in addition to reduced Bowman’s space at 40DPI. Females showed upregulated LV ERO1_α_ and 4HNE while males exhibited reduced PDI and Prdx4 paired with elevated BiP. EN460 attenuated cardiac and renal dysfunction and lowered LV ERO1_α_/4HNE in females, but not males. IRE1_α_C148S mitigated cardiac dysfunction in males and restore renal indices in both sexes.

**Conclusions:** nPTX-EAE causes clinically relevant, sex-specific cardiorenal dysfunction linked to distinct ER stress/UPR alterations. ERO1_α_ inhibition protects females, whereas enhanced IRE1_α_ activity protects males and kidneys across sexes, supporting sex-specific ER stress–targeted therapies for MS-associated cardiorenal disease.

## 1. INTRODUCTION

The prevalence and incidence of autoimmune disease is increasing worldwide by approximately 19% with women at a fourfold increased risk compared to men^1,2^. Multiple sclerosis (MS), is neurodegenerative autoimmune disorder and is the most prevalent chronic inflammatory disease of the central nervous system (CNS) affecting over 2 million people worldwide with approximately 70% of patients being women^3,4^. MS has classically defined by focal demyelination and progressive neurologic disability^5–7^. However, beyond the CNS, epidemiologic and clinical studies increasingly recognize that MS patients experience an elevated burden of cardiovascular and renal comorbidities. More specifically, the risk of heart failure is 68% greater in MS patients and 3-15% experience renal disease that contribute substantially to morbidity and mortality complicating disease management^8–13^. Despite the clinical importance, mechanisms by which neuroinflammation facilitates peripheral organ dysfunction remain poorly defined and limit the development of targeted mechanisms-based interventions.

Experimental autoimmune encephalomyelitis (EAE) is one of the most commonly used preclinical model of MS and recapitulates many pathologies found in human disease including immune cell infiltration, demyelination, and sensorimotor deficits^14,15^. While EAE has been extensively characterized for CNS pathology and behavioral outcomes, there is only one study that has evaluated cardiac physiology in EAE suggesting the presence of clinically relevant cardiac dysfunction and none investigate renal function^16^. Therefore, it is necessary to further interrogate the cardiorenal performance in EAE to define peripheral consequences of systemic neuroinflammation and identify mechanistic mediators of organ injury.

Oxidative and endoplasmic reticulum (ER) stress are central, convergent mechanisms that have been shown to link inflammation to tissue injury in both the heart and kidney^17–19^. Systemic inflammation increases production of reactive oxygen species (ROS) and lipid peroxidation impairing sarcomere function, promotes microvascular dysfunction, and drives maladaptive remodeling^20–22^. In the ER, disulfide bond formation during oxidative protein folding is a critical source of peroxide species. ER oxidoreductin-1_α_ (ERO1_α_) catalyzes the reoxidation of protein disulfide isomerase (PDI), producing H_2_O_2_ as a byproduct coupling protein folding to redox homeostasis^23^. Perioxiredoxin-4 (Prdx4) provides an alternative ER oxidative pathway and, together with PDI, mediates unfolded protein response (UPR) capacity to restore proteostasis^23,24^. Dysregulation of ERO1_α_, PDI, and/or Prdx4 can disrupt ER-mitochondrial Ca2+ exchange, exacerbate oxidative stress, and sensitize organs to cellular injury. The UPR sensor inositol-requiring enzyme 1_α_ (IRE1_α_) mediates adaptive responses to ER stress and, when selectively activated, can attenuate inflammatory signaling and cell death^25,26^. In contrast, dysregulated ERO1_α_ activity can amplify oxidative injury. These opposing roles suggest that modulated ER oxidoreductases or adaptive UPR signaling may be a strategy by which inflammation-driven cardiorenal injury is attenuated.

In this study, we performed comprehensive echocardiographic and renal Doppler assessments of both female and male EAE mice when CNS motor deficits were established. We analyzed left ventricle (LV) expression of ERO1_α_, PDI, and Prdx4, quantified lipid peroxidation, and tested whether pharmacologic ERO1_α_ inhibition (EN460) or sustained IRE1_α_ activation (C148S genetic model) ameliorates cardiorenal dysfunction. We found that EAE causes concurrent systolic and diastolic impairment and renal dysfunction in both sexes. LV ERO1_α_ and PDI/Prdx4 are dysregulated in a sex-dependent manner and ERO1_α_ inhibition and enhanced IRE1_α_ activity confer sex-specific protection against cardiorenal injury. This study is the first to identify ER oxidoreductases and UPR signaling as mechanistic mediators of EAE-induced cardiorenal dysfunction and emphasize the need the development of sex-specific therapeutic interventions.

## 2. MATERIALS AND METHODS

### 2.1 Mice

Female and male C57BL/6J mice were purchased at 10-12 weeks of age from The Jackson Laboratory (Strain #000664). IRE1_α_ C148S mice, originally generated using the CRISPR technology in the Transgenic and Chimeric Mouse facility of the Department of Genetics at University of Pennsylvania, were maintained through an in-house colony. Animals were housed in a 12/12-hour (hr) light/dark cycle room and given food and water *ad libitum*. Mice were acclimated to the colony housing room for 1 week prior to handling and experiments. All animal-use experimental protocols were performed with the approval of The George Washington University Institutional Animal Care and Use Committee (#A2024-109) and in accordance with the United States Public Health Service’s Policy on Humane Care and Use of Laboratory Animals.

### 2.2 nPTX-EAE induction and Pharmacological Treatment

Mice were immunized via a subcutaneous injection containing an emulsion of 100µg of MOG_35-55_ peptide and complete Freund’s adjuvant supplemented with 200µg of heat-inactivated Mycobacterium tuberculosis H37Ra as previously described^14,27^. Immunizations were administered 50µL each to the right and left hind flanks. A booster MOG/CFA injection was given to all mice 7 days after the first immunization. Controls mice received the same immunization components except in the absence of the MOG_35-55_ peptide. In studies investigating the inhibition of ERO1_α_ activity, EN460 (MedChemExpress, Cat #HY-12837) was intraperitoneally administered at 10mg/kg. Administration began 10 days post-immunization (DPI) and was done twice weekly until termination of experiment. Vehicle and diluent for drug delivery consisted of dimethyl sulfoxide (DMSO) and corn oil. Those performing behavioral and physiological assays were blinded to the treatment groups each animal was assigned to prevent experimenter bias.

### 2.3. Behavioral and Physiological Assays

#### Scoring of Motor Deficits

EAE locomotive clinical symptoms were assessed daily by open-feel testing using a standard scoring from 0 to 6 as previously described^14,27,28^: 0 – no motor deficits, 1 – 50% loss of tail tone, 2 – fully flaccid tail, 3 – hind-limb paralysis (<50% plantar stepping during open field observation), 4 – forelimb paralysis, 5 – moribund, 6 – death.

#### Echocardiograms

Transthoracic echocardiography was conducted on mildly sedated mice creating live images between 36-38DPI. Mice were placed in the supine position on a heated stage using on stage electrodes to monitor heart and respiration rates. Mice were anesthetized with isoflurane (1-1.5% isoflurane and 98.5-99% O2) and the temperature was maintained at 37°C. The mice chests were shaved, and Nair cream was used to remove fur. Ultrasonic gel was added to the chest region for measurement by 22 to 55 MHz echocardiography transducer (MS550D; Vevo 2100, FUJO-FILM VisualSonics). The heart rate was kept between 400 and 550 beats per minute (bpm). 2-dimensional images were obtained from the parasternal long axis view. M-mode images were obtained from the short axis view. Mitral valve flow was assessed through color doppler and tissue doppler. Left ventricular parameters were calculated by averaging values over 5 cardiac cycles.

#### Renal Ultrasounds

Abdominal ultrasounds were conducted on mildly sedated to visualize the kidneys between 36-38DPI. Mice were placed in the supine position on a heated stage with on stage electrodes monitoring heart and respiration rates. Mice were lightly anesthetized with isoflurane (1-1.5% isoflurane and 98.5-99% O2) and the temperature was maintained at 37°C. Murine heart rates were kept above 400 bpm. Ultrasonic gel was added to the abdominal region for measurement by 22 to 55 MHz echocardiography transducer (MS550D; Vevo 2100, FUJO-FILM VisualSonics). 2-dimensional images (B-mode) were obtained for long axis and short axis of both the right and left kidney. Color doppler was used to assess renal arterial flow measuring peak systolic velocity, late diastolic velocity, and renal restrictive index. Renal parameters were calculated by averaging values from 5 cycles.

### 2.4 Tissue Collection

#### Fresh Tissue

At 40DPI, mice were deeply anesthetized using ketamine/xylazine and transcardially perfused with 1x PBS. Left ventricle and kidney were extracted and placed in 1.5mL microcentrifuge tubes on dry ice. Tissues were homogenized in 1x RIPA buffer (Cell Signaling 10x RIPA, Cat #9806; 20mM Tris-HCl ph7.5, 150mM NaCl, 1mM Na2EDTA, 1mM EGTA, 1% NP-40, 1% sodium deoxycholate, 2.5mM sodium pyrophosphate, 1mM beta-glycerophosphate, 1mM Na3VO4, 1µg/mL leupeptin) supplemented with protease inhibitor (ThermoFisher, Cat #78429) and phosphatase inhibitor (Cell Signaling, Cat #5870). The samples were then centrifuged at 4°C for 15 minutes (min) at 13,200RPM. Protein concentrations from resulting supernatant were determined using the DC™ Protein Assay Kit (Bio-Rad, Cat #5000115).

#### Fixed Tissue

At 40DPI, mice were anesthetized using ketamine/xylazine and transcardially perfused with 1x PBS and then with 10% neutral buffered formalin (NBF, pH 7.4). Hearts and kidneys were dissected and post-fixed overnight at 4°C in 10% NBF. Hearts and 1 kidney from each animal were then immersed in 30% sucrose for dehydration for 48 hr at 4°C. These tissues were then embedded in Optimal Cutting Temperature compound (O.C.T., Tissue-Tek^®^) and stored at −80°C until sectioned. Hearts were sectioned at 10µm and kidneys were sectioned at 8µm. The 2^nd^ kidney from each animal was transferred to 70% ethanol (EtOH) after 24 hr in 10% NBF and then processed for paraffin embedding and sectioned at 3µm thickness using The George Washington University’s Research Pathology Core Lab.

### 2.5 Biochemical Assays

#### Western Blot

Protein samples were separated using 7.5-12% sodium dodecyl sulfate (SDS)-polyacrylamide stain-free gels and transferred to nitrocellulose membranes using the Turbo Blot system (Bio-Rad). Total protein was image using Bio-Rad’s stain free imaging system. Membranes were blocked for 1 hour in 5% bovine serum albumin (BSA). Primary antibodies were diluted in blocking buffer and incubated overnight at 4°C: ERO1_α_ (Abcam, rabbit, 1:500), PDI (Cell Signaling, rabbit, 1:500), Prdx4 (Abcam, rabbit, 1:500), 4-HNE (Abcam, rabbit, 1:750), BiP (Abcam, rabbit, 1:1500). Following primary incubation, membranes were incubated in horseradish peroxidase (HRP)- or fluorescently-conjugated secondary antibodies. Protein bands were detected using a chemiluminescent substrate for HRP-conjugated secondaries (West Pico, ThermoFisher) and imaged using Bio-Rad’s ChemiDoc system. Band intensities were quantified using ImageLab Software and proteins were normalized to total protein per sample using Bio-Rad’s stain-free total protein imaging and subsequently normalized to the control conditions for graphing quantification as previously described^14,27,29^.

#### Hematoxylin and Eosin (H&E) Staining

Slides containing sectioned hearts were first rinsed with 1x PBS for 3 min followed by a 3 min rinse of diH_2_O. Slides were then immersed in Mayer hematoxylin for 5 min followed by a 1 min wash in diH_2_O. Slides were dipped/immersed once in 0.5% acid alcohol (70% EtOH, 0.5mL HCl) and then washed 3 times in diH_2_O for 1 min each. Slides were again washed in 1x PBS for 1 min followed by another 3 washes in diH_2_O for 1 min each. Counterstaining in alcoholic eosin was done for 1 min 45 sec. Slides were then dehydrated by immersing in 3 changes of 95% EtOH, 2 changes of 100% EtOH for 1 min each, and subsequently 3 changes of Xylene 1 min each. Slides were then mounted and cover slipped using DPX (Millipore Sigma, Cat #06522-500ML).

#### Periodic Acid Schiff (PAS) Staining

Paraffin sectioned kidneys were washed twice in xylene for 4 min each followed by 2 washes in 100% EtOH 2 min each. Slides were then washed once in 95% EtOH and once in 70% EtOH for 2 min each followed by a rinse in diH_2_O. The slides were immersed in 0.5% Periodic Acid (Millipore Sigma, Cat #P7875-25MG) solution for 5 min. This was then followed by 3 rinses in diH_2_O. Slides were incubated in Schiff’s reagent (Millipore Sigma, Cat #3952016-500ML) for 15 min and rinsed with diH_2_O for 5 min. Incubation in hematoxylin (GHS216-500ML) was done for 90 seconds (sec). Rinses in diH_2_O were then performed 6 times. Following the rinses, one wash in 70% EtOH and one wash in 95% EtOH was done for 2 min followed by 2 washes in 100% EtOH for 2 min each. A final incubation in xylene was done for 5 min and slides were mounted and cover slipped using DPX.

### 2.6 Statistics

GraphPad Prism 10.5 (GraphPad 10, San Diego, CA, USA) was used to analyzed data and are presented as mean ± SEM. Onset of motor symptoms and quantification of histological samples were analyzed using an unpaired, two-tailed Mann-Whitney *t*-test (_α_ = 0.05). One-way ANOVA (_α_ = 0.05) with comparisons made between groups using Tukey’s multiple comparisons test or an unpaired, two-tailed Mann-Whitney *t*-test (_α_ = 0.05) were used to analyze western blots. Data outliers were identified and excluded using the ROUT test (Q = 1%).

## 3. RESULTS

### 3.1 EAE induces cardiac dysfunction in both sexes through impaired contractility

Previous work from our group has demonstrated that 60-80% of mice present with motor deficits following EAE induction in conjunction with lumbar spinal cord and cortical demyelination and inflammation^14,27,28,30^. Despite the robust characterization and investigation of underlying CNS pathology associated with sensory and locomotive deficits in EAE, there has been one study published^16^ that has begun to address a prevalent comorbidity experienced by MS patients in EAE which is cardiac dysfunction and heart failure^31–33^. This prompted us to determine if our EAE model developed cardiac dysfunction. To test this, we performed echocardiograms between 36-38 days post-immunization (DPI) to ensure that our mice were developing motor symptoms prior that are characteristic of CNS pathology and as an indicator of successfully induction. Similar to previous studies we have published^14,27^, 67% of EAE mice developed motor symptoms with earlier onset in females compared to males (Supplemental Fig.1A-D).

To investigate cardiac function post-EAE induction, we measured several parameters of systolic and diastolic function. Following nPTX-EAE induction, both females and males had reduced ejection fraction (EF), fractional shortening (FS), and stroke volume (SV; Fig.1A-C). Despite no significant change in the E/A ratio of peak velocity blood flow during LV relaxation in early diastole (E wave) to peak velocity in late diastole during atrial contraction (A wave); Fig. 1D), there are increases in isovolumic contraction (IVCT; Fig.1E) and relaxation times (IVRT; Fig.1F). While LV filling pressure is elevated in the males only (E/e’; Fig.1G), the Myocardial Performance Index, MPI ((IVCT+IVRT))/ejection time) shows impaired cardiac function in both sexes (Fig.1H).

**Figure 1.**
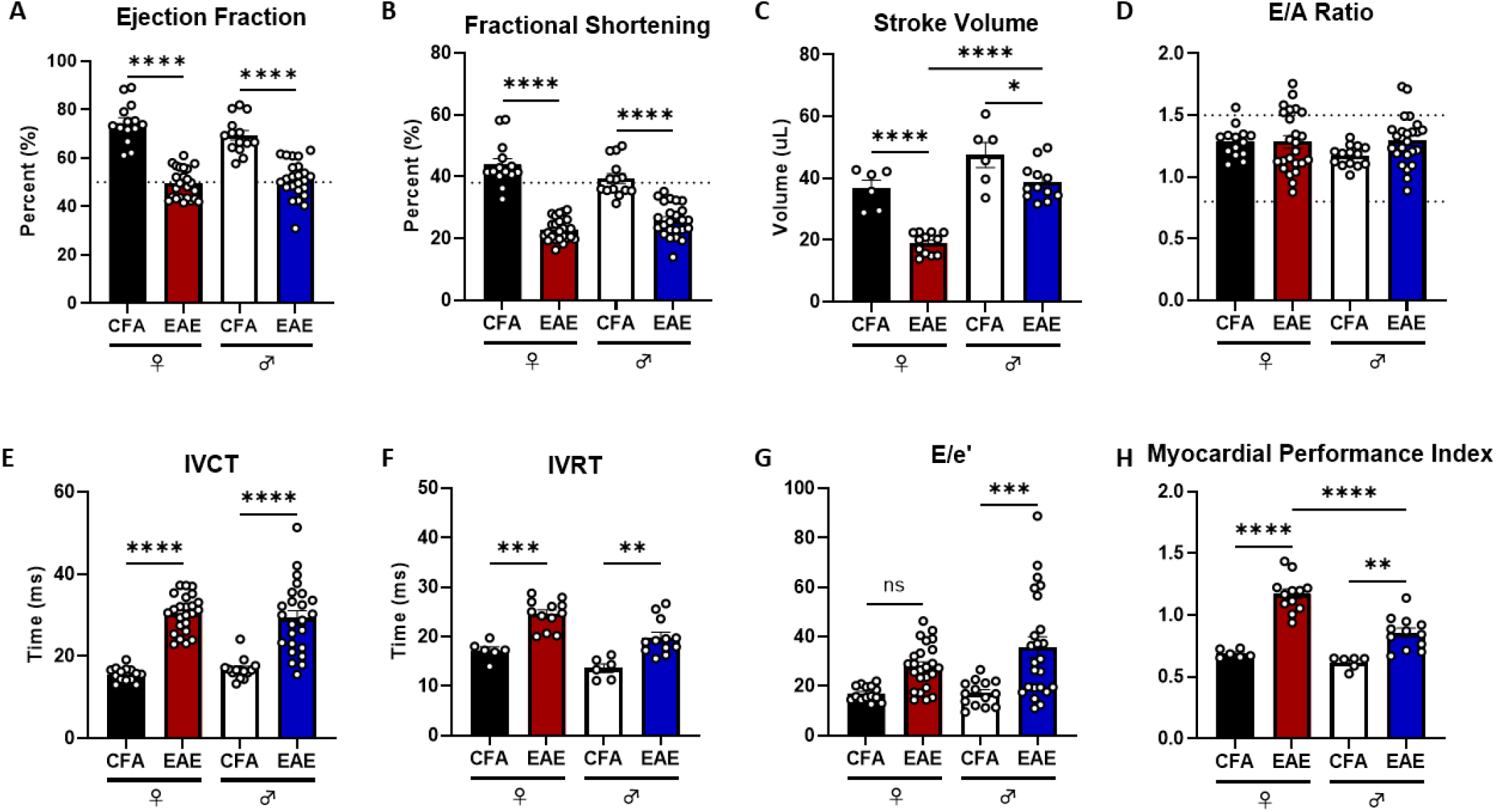
EAE causes both systolic and diastolic dysfunction in females and males that contribute to reduced cardiac performance. There is reduced (**A**) ejection fraction, (**B**) fractional shortening, (**C**) and stroke volume in females and males. (**D**) While there was no change in E/A ratio, (**E**) IVCT and (**F**) IVRT are elevated in both sexes. (**G**) E/e’ is elevated in the males only, however, the (**H**) myocardial performance index is elevated in both females in males (*p<0.05, **p<0.01, ***p<0.001, ****p<0.0001; data represented as mean ± SEM).

To determine if cardiac dysfunction was mainly due to hypertrophy or impaired relaxation-contraction mechanisms, we measured heart mass and structural integrity via hematoxylin and eosin (H&E) staining. No changes to heart mass were found nor was cardiac hypertrophy present (as calculated by dividing heart weight by tibia length cubed (HW/TL^3^ Supplemental Fig.2). H&E staining of the heart reveals reduced thickness and increased poration in the LV of females (Supplemental Fig.3A,C,E,F) and reduced thickness in the males (Supplemental Fig.3B,D,G,H).

### 3.2 EAE induces renal dysfunction in both sexes

Previous research has established a strong link between cardiovascular disease and renal dysfunction with heart failure closely linked with worsening renal function as indicated by impaired kidney filtration rates^12,34,35^. Among MS patients, between 3-15% of patients exhibit renal disease, but risk of developing both cardiac and renal dysfunction is elevated with the use of medications to treat disease or secondary complications^11,13,36–38^. In addition to cardiac dysfunction, nPTX-EAE induction leads to increased blood flow resistance in the kidney through the renal arteries (renal resistive index [RRI]; Fig.2A) and reduced blood flow through renal vessels when the heart relaxes in diastole (end diastolic velocity [EDV]; Fig.2B). Renal function in males also shows some increase in blood flow speed through the renal artery (peak systolic velocity [PSV]; Fig.2C). Increased RRI and reduced EDV is commonly associated with impaired glomerular filtration rate. A reduction in Glomerular filtration rate (GFR) is linked to increased cardiovascular risk and can also be associated with reduced Bowman’s space^39–41^. Both EAE females and males exhibit reduced Bowman’s space compared to controls at 40DPI (Supplemental Fig.4). However, there is a lack of renal hypertrophy with no change in organ size (Supplemental Fig.5).

**Figure 2.**
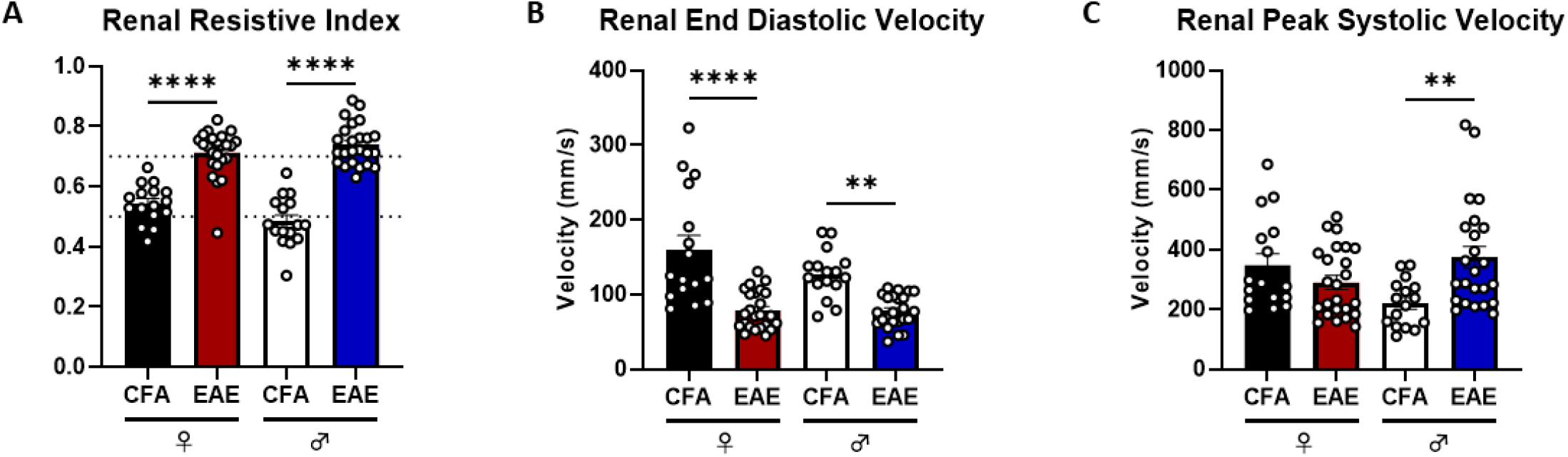
EAE causes increased blood flow resistance in renal blood vessels. (**A**) EAE females and males have increased resistance in renal blood flow along with (**B**) reduced renal end diastolic velocity. (**C**) Renal peak systolic velocity is elevated in the males only (**p<0.01, ****p<0.0001; data represented as mean ± SEM).

### 3.3 Biologically distinct LV protein expression of oxidative stress and UPR markers in EAE

Inflammation is known to promote generation of reactive oxygen species (ROS) and disrupt unfolded protein response (UPR), which then further drives tissue injury and inflammatory signaling^17^. A critical endoplasmic reticulum (ER) enzyme named ER oxidoreductin 1 alpha (ERO1_α_) produces ROS, regulates Ca^2+^ release, and controls ER-mitochondrial communication^19^. Hypoxic conditions, which are triggered in response to inflammation and associated with MS, have previously been shown to induce transcription of hypoxia-inducible factor 1 (Hif1_α_) that then increases ERO1_α_ expression^42,43^. Dysregulated ERO1_α_ activity impairs Ca^2+^ exchange, redox balance, and enhances ROS production as indices of environmental oxidative stress. ERO1_α_ produces ROS specifically in the form of H_2_O_2_ as a byproduct from oxidative protein folding in the ER. H_2_O_2_ results in lipid peroxidation that can be measured through quantification of 4-hydroxynonenal (4-HNE)^44,45^. Following EAE induction there is an increase in ERO1_α_, 4HNE, and Hif1_α_ expression in the female LV (Fig.3A-C). There is no change in expression of proteins in female kidneys (Supplemental Fig.6A-C). However, there is no change in the male LV for ERO1_α_ and Hif1_α_, but there is a reduction in 4HNE (Fig.3D-F). Similarly, in the male kidneys there is no change in ERO1_α_ or Hif1_α_ expression; however, there is an increase in 4HNE (Supplemental Fig.6D-F).

**Figure 3.**
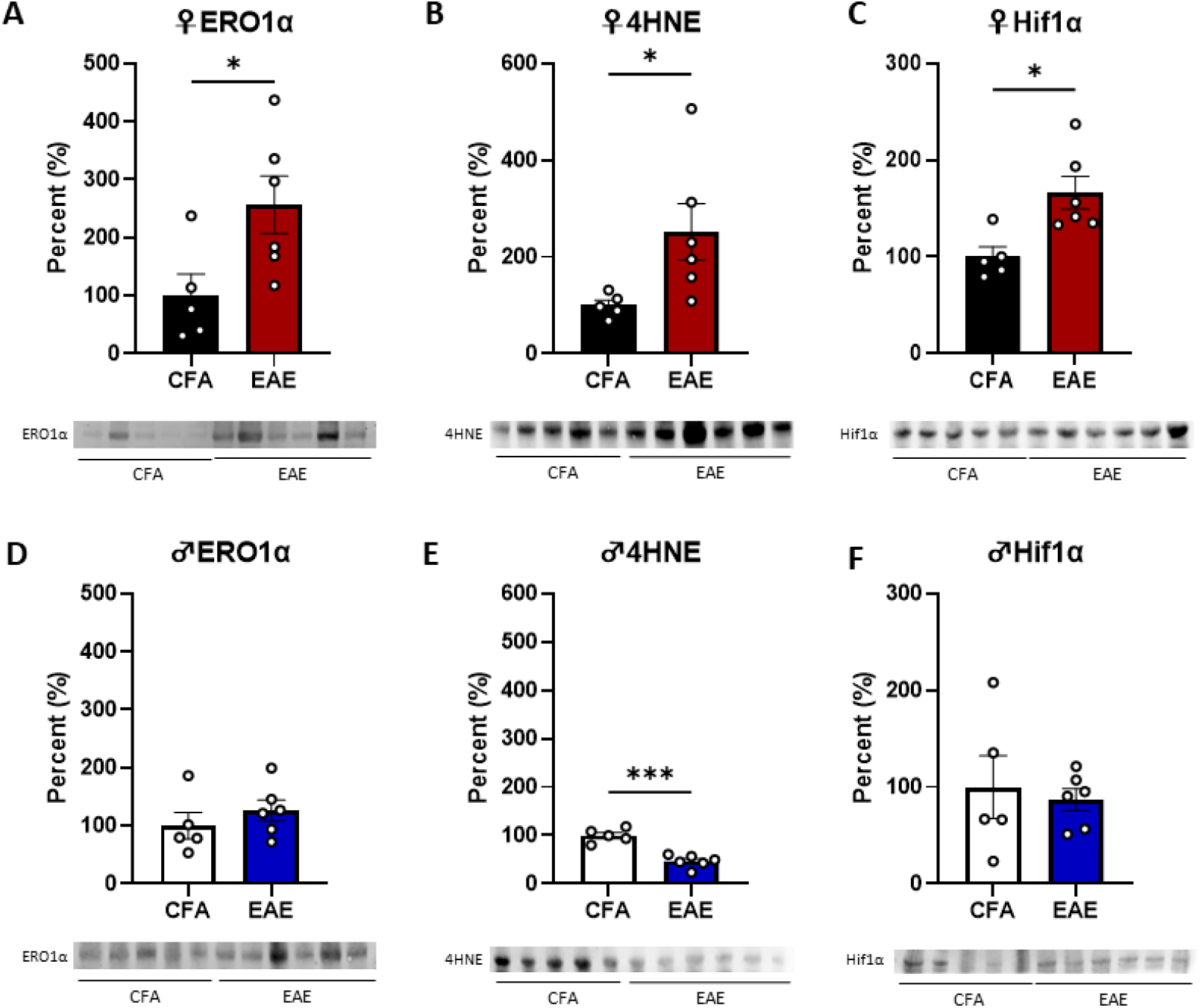
Proteins associated with the oxidative stress are elevated in the female left ventricle. Expression of (**A**) ERO1, (**B**) 4HNE, and (**C**) Hif1_α_ are elevated in female LV. In males, there is no change in (D) ERO1, (E) reduced 4HNE, and (F) no change in Hif1_α_ (*p,0.05, ***p<0.001; data represented as mean ± SEM).

ERO1_α_ has also been shown to participate in the UPR through interactions with protein disulfide isomerase A1 (PDIA1) facilitating the formation of disulfide bonds for protein folding and binding immunoglobulin protein (BiP) acting as chaperone binding to unfolded protein^46,47^. However, an alternative PDIA1 oxidation enzyme that facilitates protein folding is peroxiredoxin 4 (Prdx4) and its expression in the ER is equivalent to PDI^48^. Changes in PDIA1 that mirror changes in ERO1_α_ and/or Prdx4 would suggest an impaired UPR pathway. There are no changes in PDIA1, Prdx4, or BiP expression the EAE female LV (Fig.4A-C). In the kidneys, there is no change to PDIA1 or Prdx4, but there is an increase in BiP (Supplemental Fig.7A-C). In males we observe reduced LV expression of both PDIA1 and Prdx4 with elevated BiP expression (Fig.4D-F). In male kidney, there is no change to PDIA1 or Prdx4, but there is reduced BiP expression (Supplemental Fig.7D-F).

**Figure 4.**
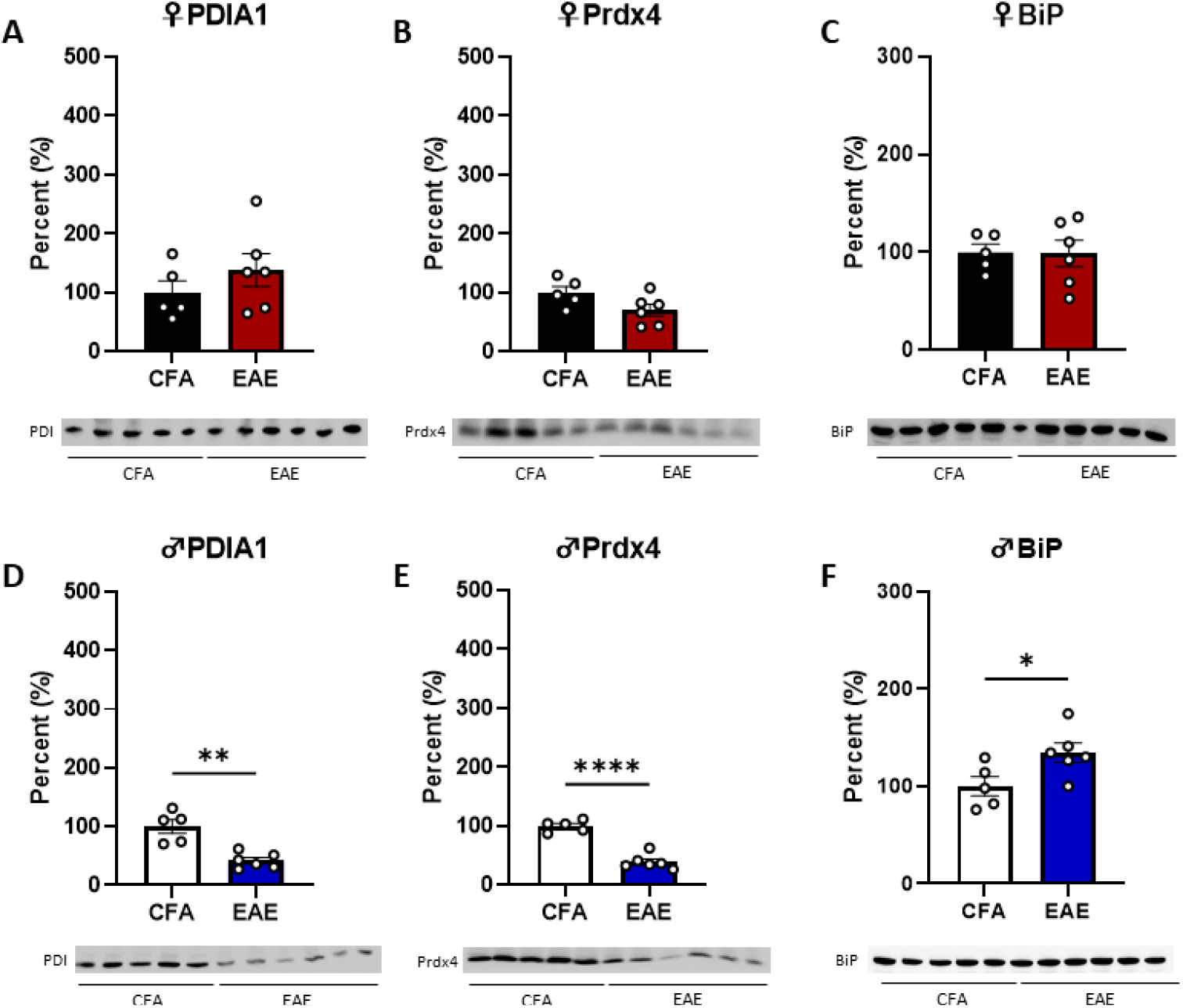
Proteins associated with UPR pathways are dysregulated in male left ventricle tissue. There is no change in (**A**) PDIA1, (**B**) Prdx4, (**C**) and BiP in female LV. In males, (**D**) PDIA1 and (**E**) are reduced with an increase in (**F**) BiP (*p,0.05, **p<0.01, ****p<0.0001; data represented as mean ± SEM).

### 3.4 ERO1 inhibition attenuates EAE cardiorenal dysfunction, lipid peroxidation, and ERO1_α_ expression in females

EN460 is an ERO1_α_ inhibitor that interacts with its active form preventing reoxidation. At a dose of 10mg/kg, EN460 was administered via intraperitoneal injection starting at 10DPI and given twice weekly. Abnormal values ejection fraction, fractional shortening, stroke volume, IVCT, E/e’, and myocardial performance index were attenuated in females treated with EN460 with no significant differences to controls (Fig.5A-F). However, there was no improvement for these cardiac parameters in males (Fig.5G-L). Aberrant changes to renal resistive index, end diastolic velocity, and peak systolic velocity were also mitigated in females, but not males (Fig.6A-F).

**Figure 5.**
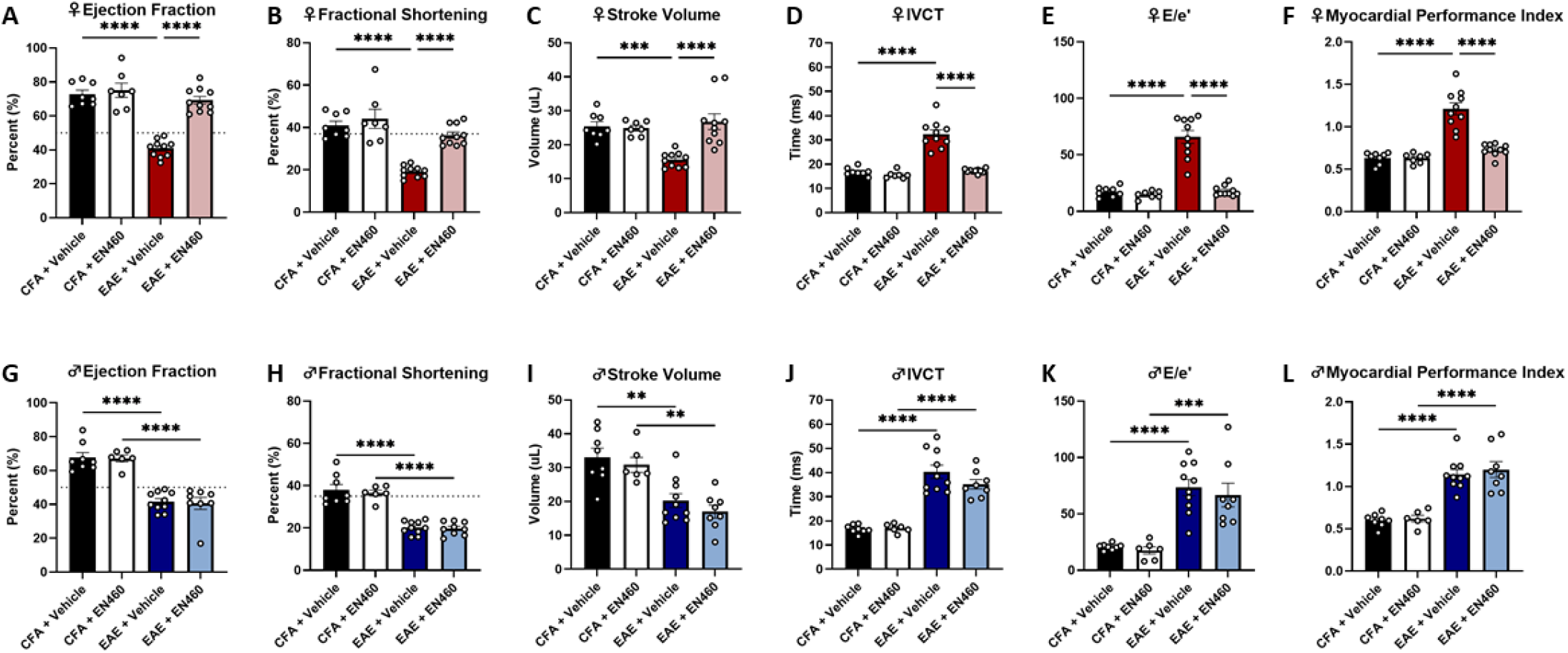
ERO1_α_ inhibition mitigates cardiac dysfunction in females, but not males. EN460 significantly attenuates impaired (**A**) ejection fraction, (**B**) fractional shortening, (**C**) stroke volume, (**D**) IVCT, (**E**) E/e’, and improves (**F**) myocardial performance index. Males treated with EN460 have no significant improvement compared to vehicle-treated controls for (**G**) ejection fraction, (**H**) fractional shortening, (**I**) stroke volume, (**J**) IVCT, (**K**) E/e’, and (**L**) myocardial performance index (**p<0.01, ***p<0.001, ****p<0.0001; data represented as mean ± SEM).

**Figure 6.**
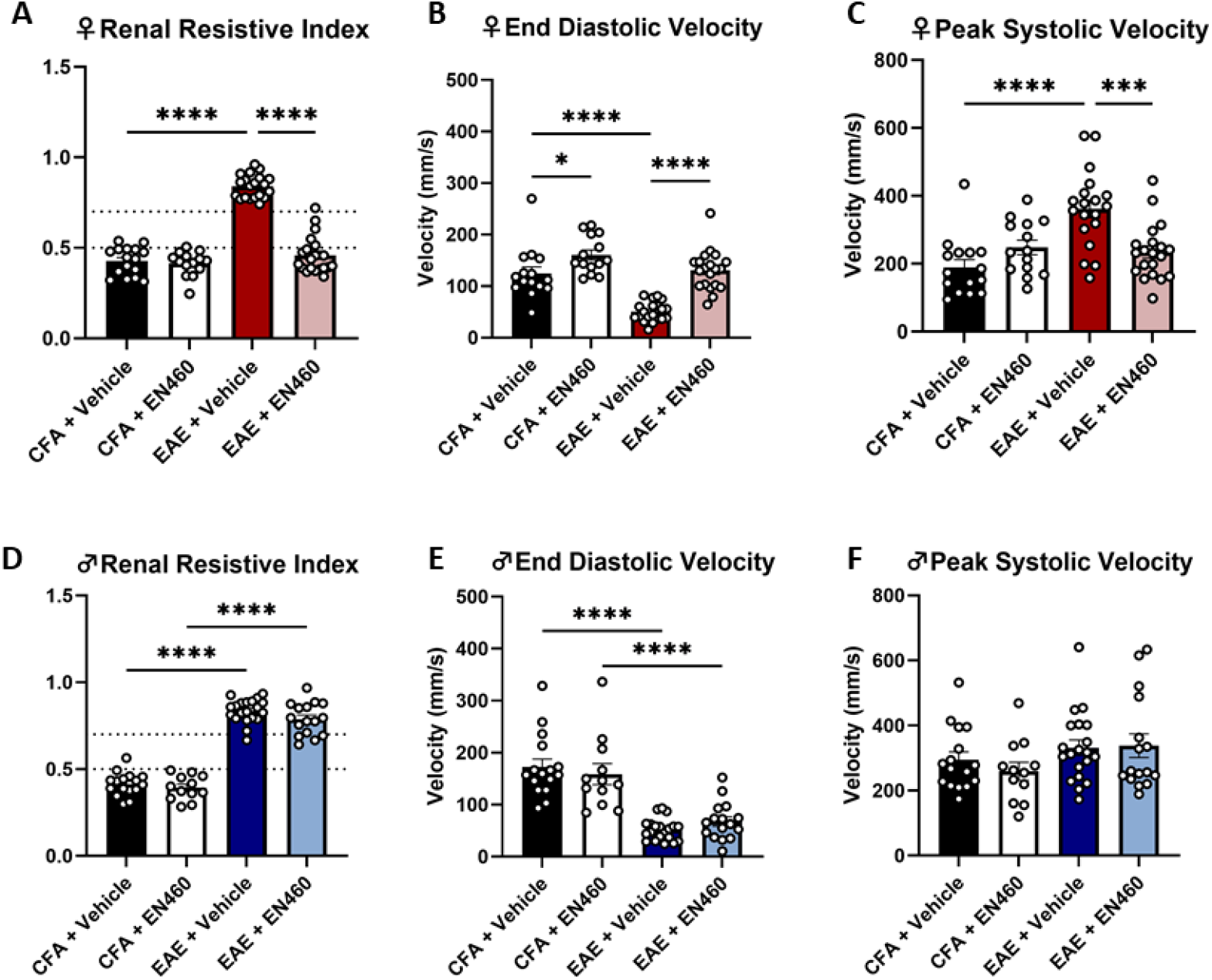
ERO1_α_ inhibition improves blood flow resistance in renal vessels in females, but not males. EN460 significantly improves (**A**) renal resistive index, (**B**) end diastolic velocity, and (**C**) peak systolic velocity in females compared to vehicle-treated EAE controls. In males, there is no change in (**D**) renal resistive index, (**E**) end diastolic velocity, and (**F**) peak systolic velocity in compared to vehicle-treated controls (*p<0.05, ***p<0.001, ****p<0.0001; data represented as mean ± SEM).

In the EN460 treated EAE females, there is reduced expression of ERO1_α_ and 4HNE with no changes to Hif1_α_ in the LV (Fig.7A-C). However, there are no changes in male LV for expression of the aforementioned proteins irrespective of treatment group (Fig.7D-F). In the kidneys of EN460 treated EAE mice, there was a reduction only in female 4HNE with no change in ERO1_α_ or Hif1_α_ (Fig.8A-C). Similar to the LV, there was no change in EN460 treated EAE males for kidney expression of ERO1_α_, 4HNE, or Hif1_α_ (Fig.8D-F).

**Figure 7.**
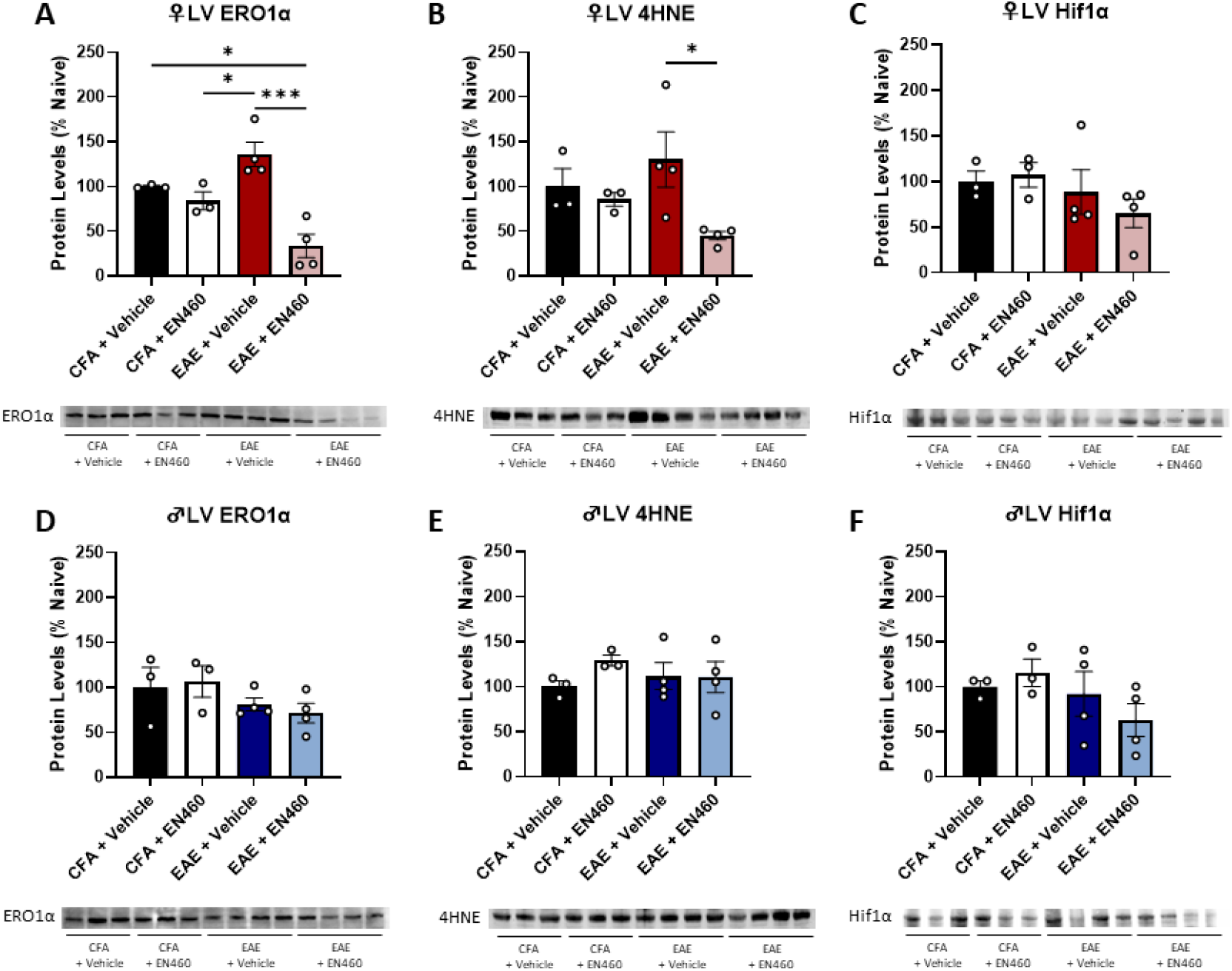
EN460 reduces EAE-induced lipid peroxidation following EAE in female left ventricle tissue. **In females**, (**A**) ERO1_α_ and (**B**) 4HNE are reduced in EN460 treated EAE females with no change in (**C**) Hif1_α_. In males, there is no change in (**D**) ERO1_α_, (**E**) 4HNE, and (**F**) Hif1_α_ (*p<0.05, ***p<0.001; data represented as mean ± SEM).

**Figure 8.**
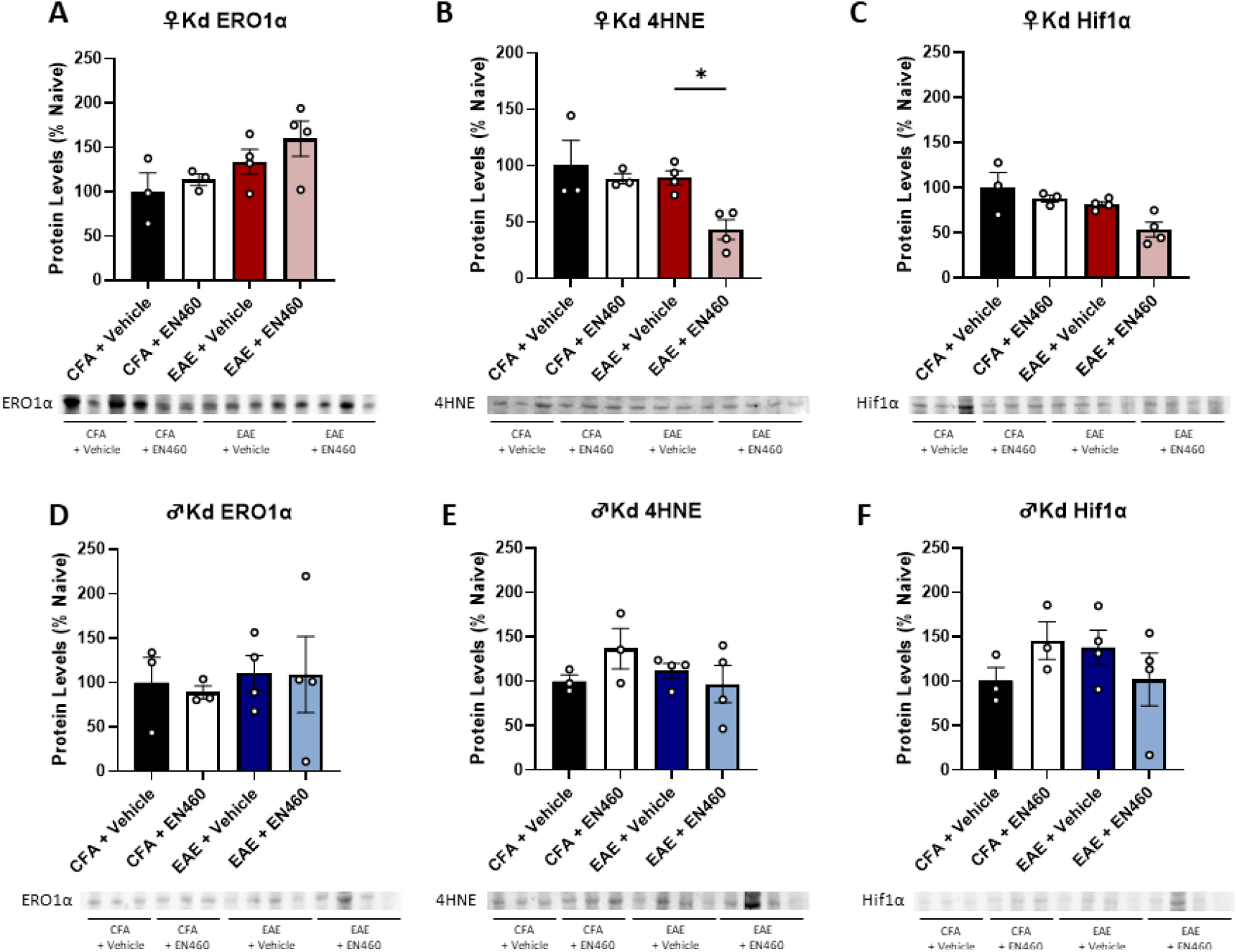
Lipid peroxidation is reduced in female EAE kidneys following EN460 treatment. In female kidneys, there is (**A**) no change in ERO1_α_, but there is a reduction in (**B**) 4HNE following EN460 treatment. (**C**) There is no change in female EAE kidney Hif1_α_. There is no change in EAE male kidneys for (**D**) ERO1_α_, (**E**) 4HNE, and (**F**) Hif1_α_ (*p<0.05; data represented as mean ± SEM).

### 3.5 Enhanced IRE1_α_ activity attenuates EAE cardiac dysfunction in males and renal dysfunction in both sexes

IRE1_α_ is an adaptive pathway activated during the UPR and is a key ER stress sensor protein. Selective activation of this pathway can be protective under stress conditions and can attenuate EAE locomotive impairment^25,26^. Since males demonstrate reduced PDI and Prdx4 that may indicate impaired UPR processes, we wanted to investigate if increased and sustained activation of IRE1_α_ mitigates cardiorenal dysfunction using a mouse model expressing a C148S variant of IRE1_α_. EAE C148S females still present with significantly altered ejection fraction, fractional shortening, stroke volume, IVCT, E/e’, and myocardial performance index compared to wild-type (WT) controls (Fig.9A-F). However, these cardiac parameters are attenuated in males compared to EAE WT controls and no significant differences to WT controls for ejection fraction, stroke volume, E/e’, and myocardial performance index (Fig.9G-L). In contrast the renal restrictive index, end diastolic velocity, and peak systolic velocity for EAE C148S show no significant difference from WT mice and dysfunction is attenuated in both sexes (Fig.10A-F).

**Figure 9.**
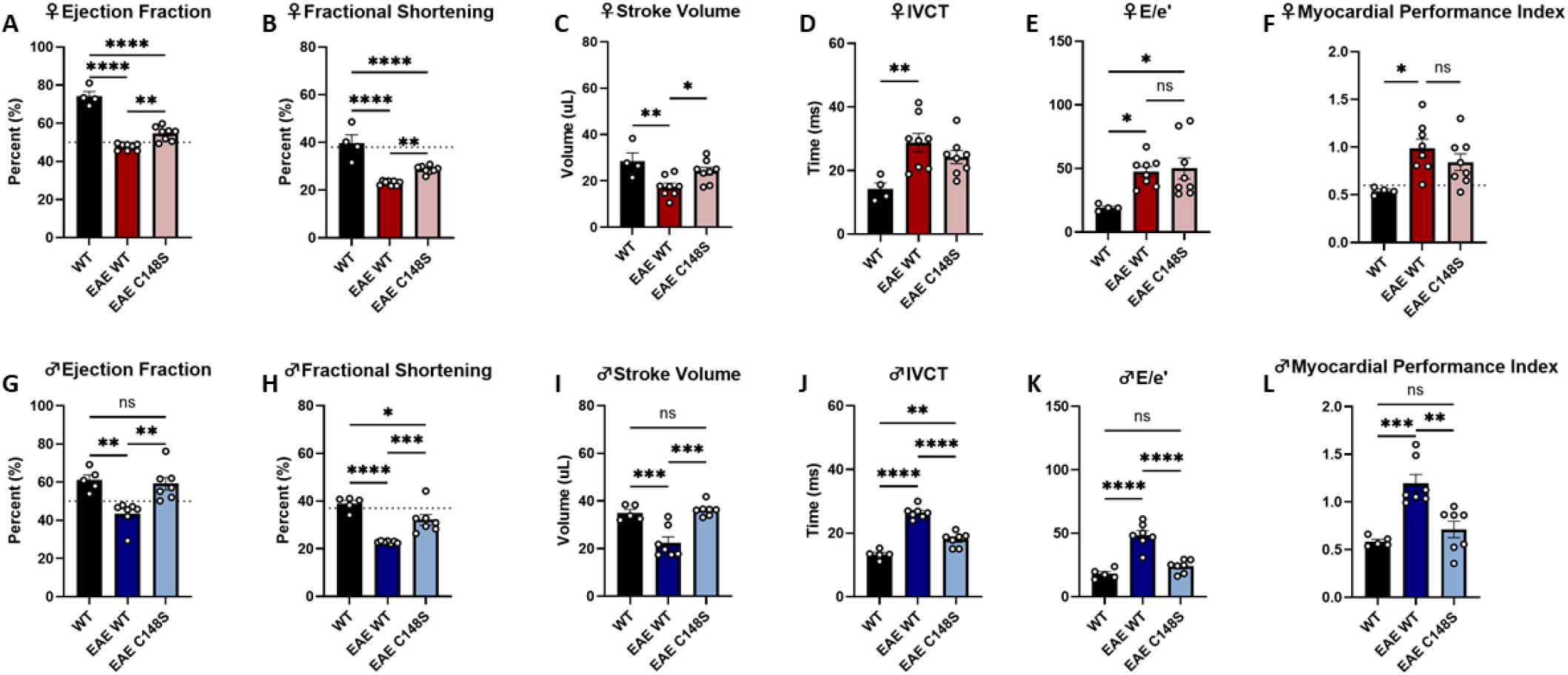
Constitutively active IRE1_α_ significantly attenuates EAE-induced cardiac dysfunction in males. (**A**) Ejection fraction and (**B**) fractional shortening are significantly reduced in EAE C148S mice compared to controls. There is a slight increase in (**C**) stroke volume and no difference in (**D**) IVCT between female C148S EAE mice and EAE WT controls. (**E**) E/e’ remains significantly elevated and there is (**F**) no difference between C148S EAE mice and EAE WT controls for myocardial performance index. C148S EAE males compared to EAE WT controls have significantly improved (**G**) ejection fraction, (**H**) fractional shortening, (**I**) stroke volume, (**J**) IVCT, (**K**) E/e’, and (**L**) myocardial performance index (*p<0.05, **p<0.01, ***p<0.001, ****p<0.0001; data represented as mean ± SEM).

**Figure 10.**
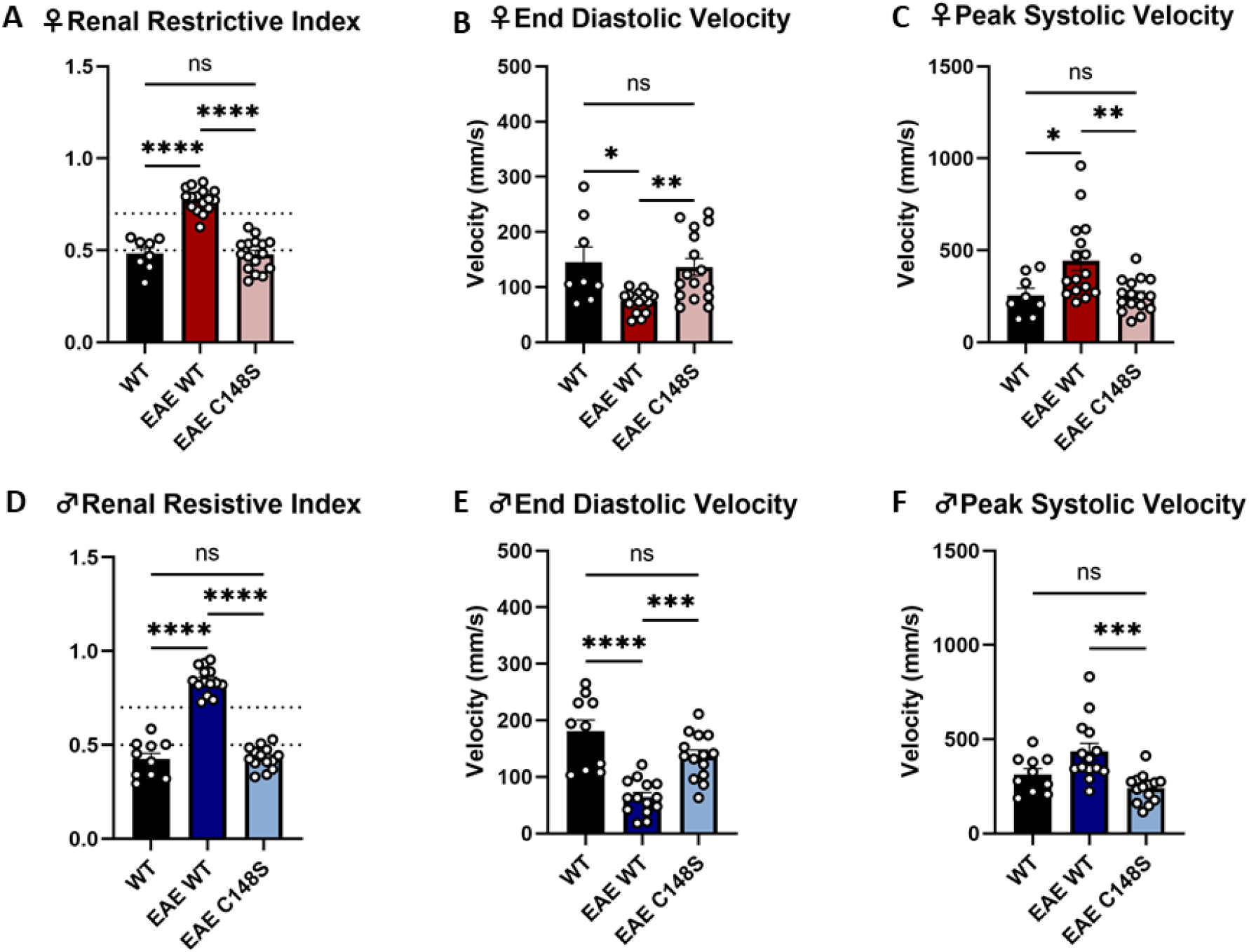
IRE1_α_ activation mitigates EAE-induced renal dysfunction in both sexes. EAE C148S females and males have significantly improved (**A**,**D**) renal restrictive index, (**B**,**E**) end diastolic velocity, (**C**,**F**) and peak systolic velocity with no difference compared to WT controls (*p<0.05, **p<0.01, ***p<0.001, ****p<0.0001; data represented as mean ± SEM).

### 3.6 Enhanced expression of UPR mediators is both sex and tissue specific

BiP, also known as GRP78, is a critical ER chaperone protein that participates in ER protein quality control. More specifically, it facilitates the folding and assembly of protein structures, targets misfolded proteins for degradation, acts as an ER stress sensor, and participates in the regulation of calcium homeostasis^49–52^. PDI is another essential ER chaperone that is induced during ER stress aiding in the formation of disulfide bonds for protein folding^53^. In the C148S EAE female LV, there was no effect on PDI or Prdx4 expression; however, there is an increase in BiP compare to WT naïve controls (Fig.11A-C). In C148S EAE males, there was increased expression of PDI and no changes in Prdx4 or BiP (Fig.11D-F). There were no changes in expression of these proteins in the female kidney and, while there is no change in males for PDI or Prdx4 following treatment, there is increased BiP expression (Fig.11G-L).

**Figure 11.**
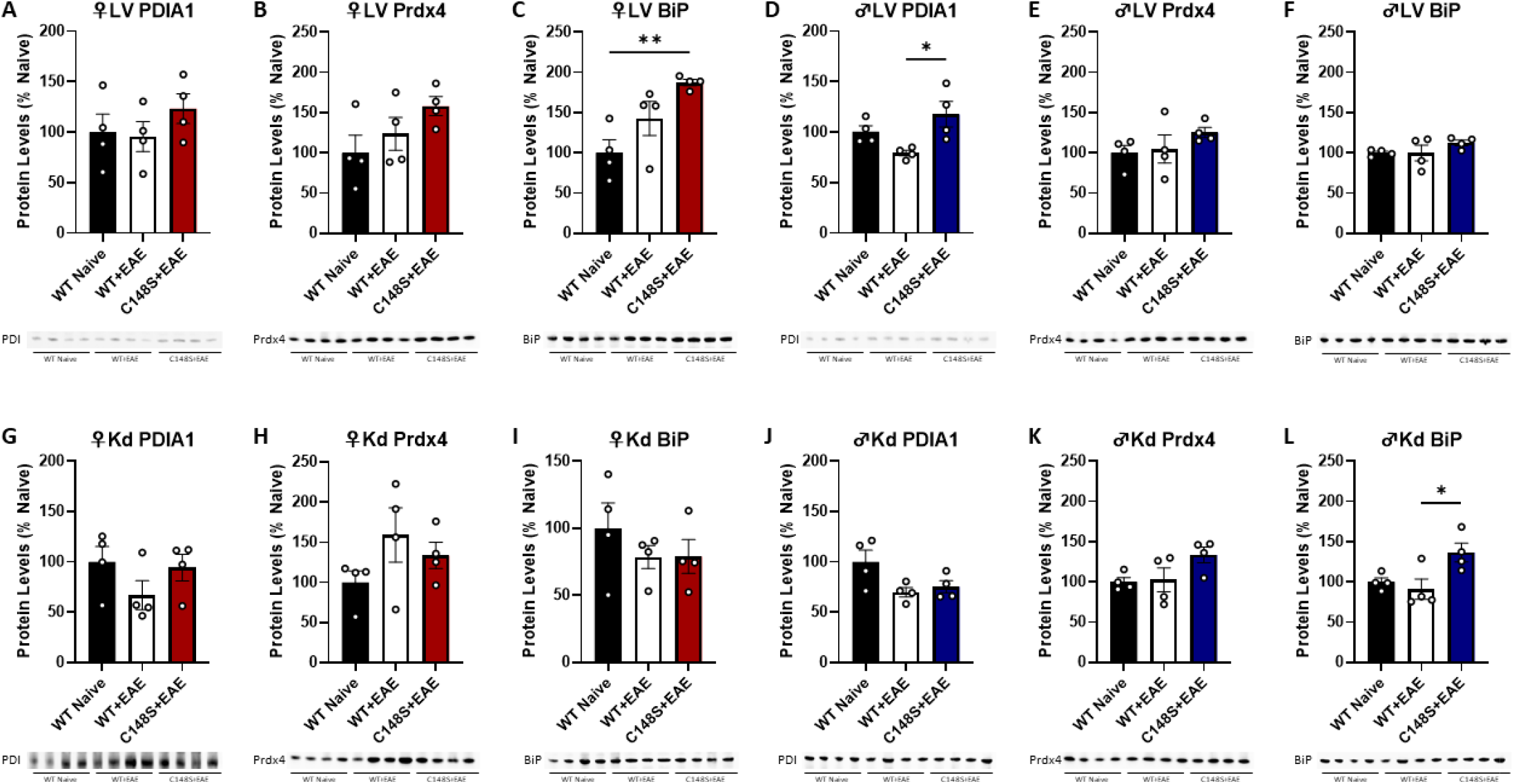
Constitutively active IRE1_α_ increases UPR activity in EAE male left ventricle and kidney tissues. In females, there is no change in LV (**A**) PDIA1 and (**B**) Prdx4. (**C**) While there is a significant increase in BiP between WT naïve and C148S EAE female LV there is no significant difference between WT EAE and C148S EAE female LV. In the male LV, there is increased (**D**) PDIA1 in C148S EAE compared to WT EAE. There is no change male LV (**E**) Prdx4 and (**F**) BiP. In female EAE kidney, there is no change in (**G**) PDIA1, (**H**) Prdx4, and (**I**) BiP. In male EAE kidneys, there is no change in (**J**) PDIA1 and (**K**) Prdx4, but there is an increase in (**L**) BiP for C148S EAE compared to WT EAE males (*p<0.05, **p<0.01; data represented as mean ± SEM).

## 4. DISCUSSION

This study demonstrates experimental autoimmune encephalomyelitis (EAE), a model of multiple sclerosis (MS), leads to significant cardiac and renal dysfunction in both sexes. These findings extend the pathophysiological impact of EAE beyond the central nervous system (CNS) and highlights sexually dimorphic mechanisms underlying peripheral organ injury. Our work reveals that cardiac and renal deficits are mechanistically linked to distinct alterations in endoplasmic reticulum (ER) stress and unfolded protein response (UPR) pathways and can be selectively attenuated through targeted intervention.

### 4.1 EAE induces systolic and diastolic cardiac dysfunction independent of structural remodeling

Oxidative stress, chronic inflammation, and altered cardiomyocyte structure have been linked to cardiovascular dysfunction in MS patients^54^. Impaired systolic and diastolic function and cardiomyopathy are increasingly recognized in MS patients, especially in the absence of overt cardiovascular risk factors^54,55^.Our echocardiographic data indicates that both systolic (reduced ejection fraction, fractional shortening, and stroke volume) and diastolic (increased IVCT, IVRT, and myocardial performance index) dysfunction in the absence of hypertrophy or gross anatomical changes occurs in EAE. While LV filling pressure (E/e’) was selectively elevated in the males, myocardial performance was impaired in both sexes. Further histological analysis confirmed thinning of the LV wall without hypertrophic compensation, suggestive of contractile dysfunction rather than overt structural remodeling.

### 4.2 Renal dysfunction recapitulates features of cardiorenal syndrome

The observed cardiac phenotype involving impairment of both systolic and diastolic in the absence of overt hypertrophy raised the question of whether EAE could drive downstream renal dysfunction via system inflammation and potential altered hemodynamics. When investigating renal perfusion and filtration, we found that EAE mice exhibit elevated renal resistive index (RRI) and reduced end-diastolic velocity (EDV) which is consistent with impaired perfusion and glomerular filtration. A reduction in Bowman’s space in the absence of renal hypertrophy further supports a functional decline in renal filtration capacity rather than structural renal injury. These findings align with clinical reports that MS patients exhibit elevated risk for renal dysfunction, particularly under conditions of systemic inflammation or drug exposure. The presence of dual organ dysfunction supports the concept of a neuroinflammatory driven cardiorenal syndrome, wherein CNS inflammation serves a systemic driver of cardiovascular and renal impairment.

### 4.3 Sexually dimorphic ER stress pathways dysfunction

The association between upregulated ERO1_α_ expression and several disease states has been well established. ERO1_α_ facilitates production of ROS, regulates Ca^2+^ release, and controls ER-mitochondrial communication^19^. Dysregulation of ERO1_α_ activity can cause impaired Ca^2+^ exchange and redox balance, which leads to ER stress, defective unfolded protein response resolution, and cell death^56,57^. More specifically, elevated ERO1_α_ activity has been linked to cancer progression, arterial thrombosis, ischemic stroke, and regulates cardiomyocyte excitation-coupled calcium release^46,58,59^. Gene knockdown of ERO1_α_ shows reduced cancer signaling proteins and ERO1_α_ knockout mice demonstrate partial protection from progressive heart failure^46,59^.

Mechanistically, we identify sex-specific alterations in ER oxidative stress and UPR components within the LV. In females, ERO1_α_ was selectively upregulated, whereas males exhibited downregulation of PDI and Prdx4, both of which are required for oxidative protein folding and redox homeostasis. Furthermore, this is supported by females exhibiting elevated indices of hypoxia and lipid peroxidation while males may have increased chaperone requirements for unfolded protein accumulation. These biologically distinct changes suggest sex-specific vulnerabilities in ER stress pathways with oxidative burden more prevalent in females and impaired adaptive UPR in males. These findings are of particular interest given that growing recognition of ER stress and UPR dysfunction as central mediators of cardiac pathology, including diastolic dysfunction, mitochondrial instability, and inflammatory signaling. The sex-specific regulation of these pathways may offer new insight into sex-based disparities observed in cardiovascular disease prevalence and outcomes.

### 4.4 Differential roles of ERO1_α_ inhibition and IRE1_α_ activation in sex-specific cardiorenal protection

Pharmacological inhibition of ERO1_α_ with EN460 reveals a striking female-specific protection against EAE-induced cardiorenal dysfunction. EN460 normalized both systolic and diastolic cardiac performance while also retaining physiological renal hemodynamics in females. These changes coincided with reduced cardiac ERO1_α_ and 4HNE expression and reduced renal 4HNE. Together, these findings implicate oxidative protein folding and lipid peroxidation as key drivers of female cardiorenal pathology and demonstrates that the targeted inhibition of ERO1_α_ is sufficient to restore physiological function. In contrast, males exhibited no benefit from ERO1_α_ inhibition and showed no changes in functional or molecular markers of cardiorenal pathology attenuation further emphasizing the sex-dependent effect of ERO1_α_-mediated stress pathways in disease progression.

Comparatively, constitutive activation of IRE1_α_ (C148S variant) provided sex-dependent protection. Male hearts exhibited preserved systolic and diastolic cardiac function while renal dysfunction was attenuated in both sexes. Mechanistically, this protection was associated with sex- and tissue-specific engagement of UPR mediators. BiP induction was restricted to female hearts and male kidneys and PDIA1 induction was elevated in the male hearts. These changes suggest that males may compensate via PDIA1-driven oxidative folding paired with BiP mediated UPR assistance. Together, these data establish that ERO1_α_ inhibition protects females by reducing oxidative burden, while IRE1_α_ activation rescues males by potentially restoring UPR signaling, with renal benefits in both sexes. These highlight the need for and importance of precision medical treatment that accounts for sex- and tissue-specific ER stress pathways in cardiovascular and renal disease.

## Conclusion

Our findings demonstrate that EAE induces cardiorenal dysfunction in both sexes through distinct, sex-specific molecular mechanisms linked to ER stress and maladaptive UPR signaling. In females, ERO1_α_ drive oxidative injury; however, in males, impaired adaptive UPR is critical to underlying pathology. Targeted modulation of these pathways, through ERO1_α_ inhibition or IRE1_α_ activation, rescues organ function in a sex-dependent manner establishing a framework for targeted therapeutic approaches that account for biologically distinct pathological mechanisms. These findings highlight the systemic impact of neuroinflammation and identify therapeutic targets for MS-associated cardiovascular and renal complications. Our work emphasizes the translational potential of targeting ER stress pathways with sex-specificity for treating cardiovascular and renal dysfunction in MS and other autoimmune diseases.

