## Supplemental Figures for "ER oxidoreductin-1_α_ and unfolded protein response as sex-dependent drivers of cardiorenal dysfunction in experimental autoimmune encephalomyelitis"

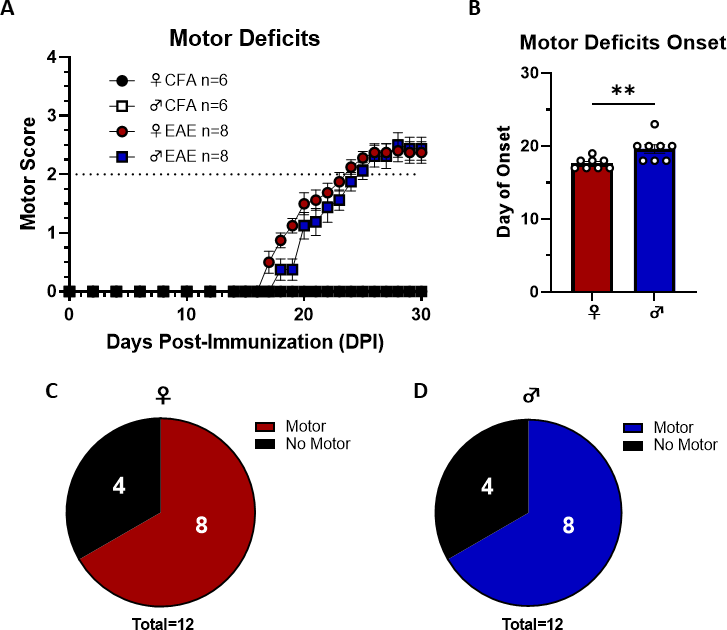

**Supplemental Figure 1: Motor deficits occur in both sexes following EAE induction with a later day of onset in males.** (**A**) EAE females and males develop motor deficits. (**B**) Onset of motor deficits are later in males than females. (**C,D**) Females and males have approximately 67% penetrance of motor symptoms (**p<0.01; data represented as mean ± SEM).

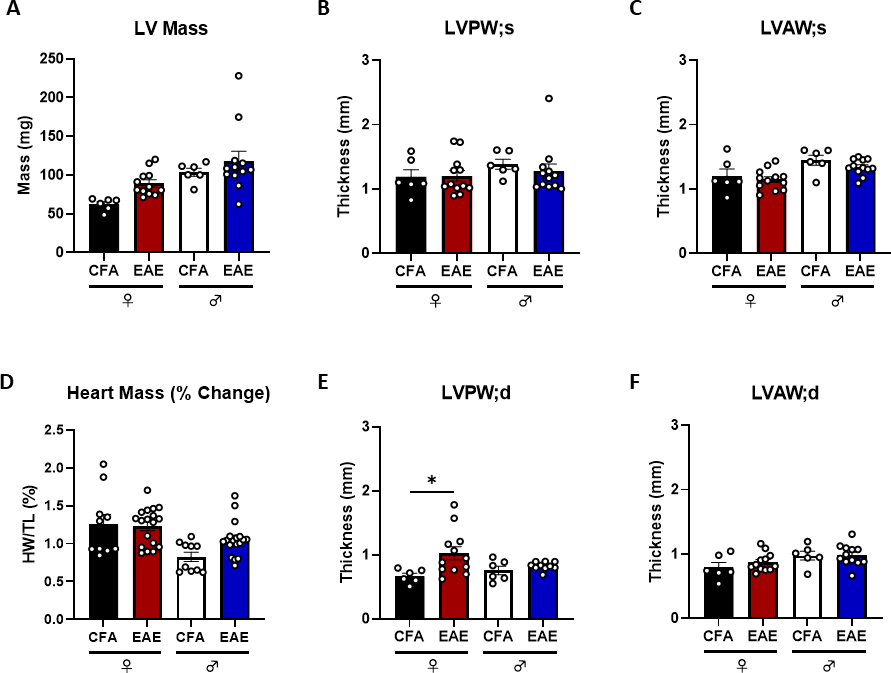

**Supplemental Figure 2: EAE does not cause significant changes to heart mass in either sex.** (**A-D**) There are no significant changes to heart mass following EAE in either sex, except for modest elevation in LVPW;d in females and (**F**) no change in LVAW;d in females or males. (*p<0.05; data represented as mean ± SEM).

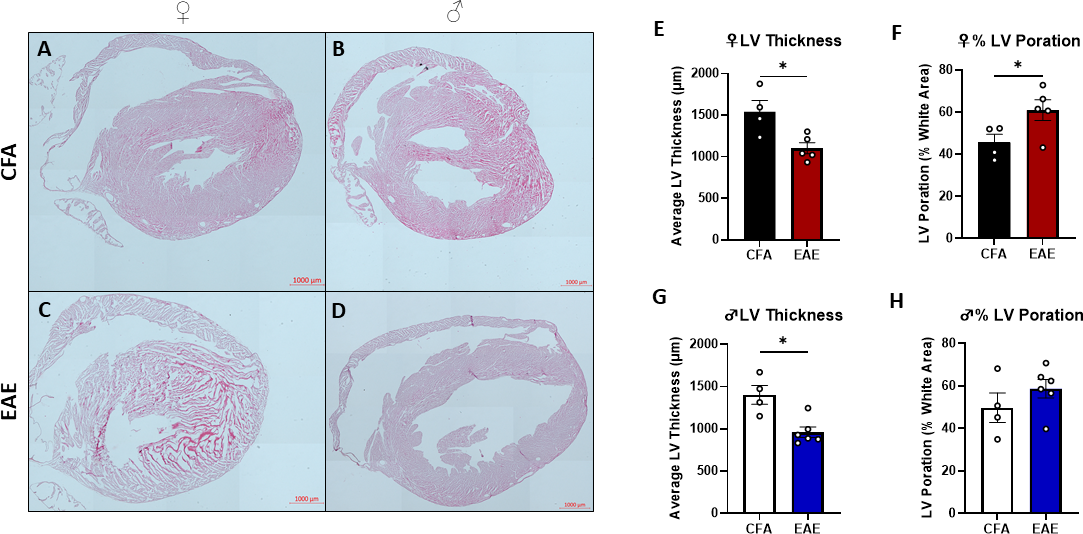

**Supplemental Figure 3: EAE causes both systolic and diastolic dysfunction in females and males that contribute to reduced cardiac performance.** There is reduced (**A**) ejection fraction, (**B**) fractional shortening, (**C**) and stroke volume in females and males. (**D**) IVCT is elevated in both sexes, with no change in (**E**) E/A ratio. (**F**) While, E/e’ is only significantly elevated in males, (**G**) IVRT is elevated in both sexes. Altogether, the (**H**) myocardial performance index is elevated in both females in males (*p<0.05, **p<0.01, ***p<0.001, ****p<0.0001; data represented as mean ± SEM).

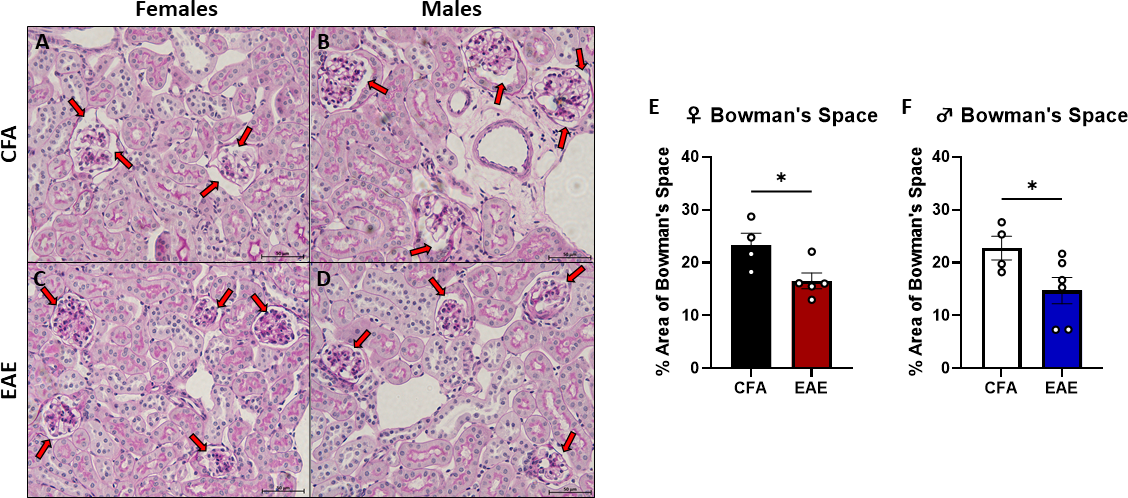

**Supplemental Figure 4: EAE causes a reduction in Bowman’s space surrounding both female and male renal glomeruli.** There is reduced Bowman’s space surrounding the glomeruli of both (**A,C,E**) females and (**B,D,F**) males (*p< 0.05; data represented as mean ± SEM).

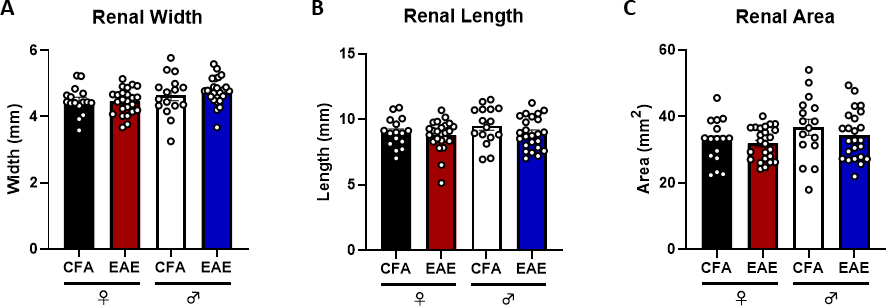

**Supplemental Figure 5: EAE does not cause changes in renal size.** (**A-D**) There are no significant changes to heart mass following EAE in either sex, except for modest elevation in LVPW;d in females and (**F**) no change in LVAW;d in females or males. (*p<0.05; data represented as mean ± SEM).

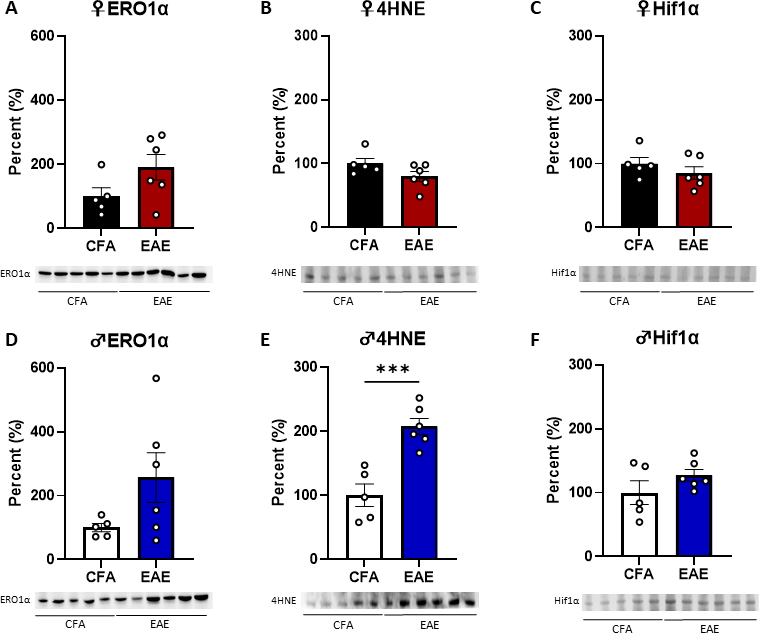

**Supplemental Figure 6. Male EAE kidneys have increased lipid peroxidation.** In EAE female kidneys, there is no change in (**A**) ERO1α, (**B**) 4HNE, and (**C**) Hif1α**.** In EAE male kidneys, there is no change in (**D**) ERO1α, there is an increase in (**E**) 4HNE, and there is no change in (**F**) Hif1α (***p<0.001; data represented as mean ± SEM).

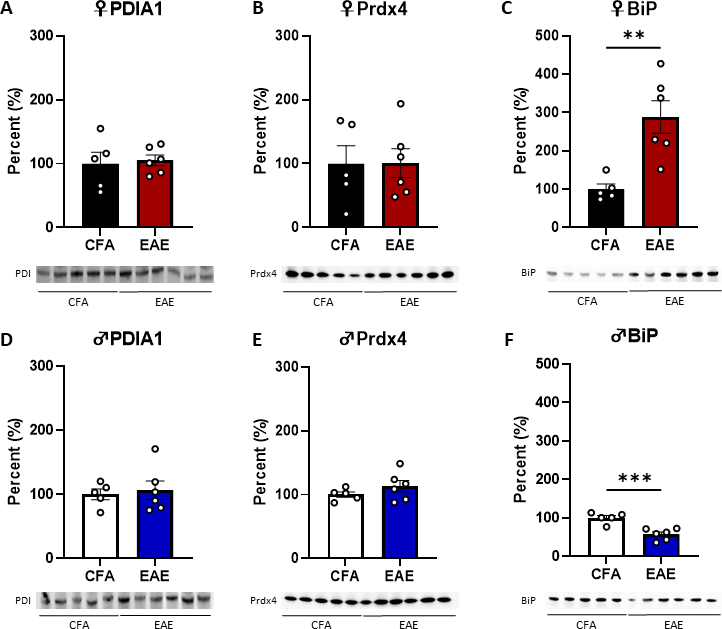

**Supplemental Figure 7. Protein chaperone BiP has biologically distinct expression in the kidney following EAE induction.** (**A**) PDIA1 and (**B**) Prdx4 remain unchanged in female EAE kidney, but there is an increase in (**C**) BiP. In males, there is also no change in EAE kidney expression of (**D**) PDIA1 and (**E**) Prdx4, but there is a reduction in (**F**) BiP (**p<0.01, ***p<0.001; data represented as mean ± SEM).
